# The effect of handrail cross-sectional design and age on applied handrail forces during reach-to-grasp balance reactions

**DOI:** 10.1101/2021.01.14.426667

**Authors:** Philippa Gosine, Vicki Komisar, Alison C. Novak

**Affiliations:** KITE Research Institute, Toronto Rehabilitation Institute – University Health Network, 550 University Avenue – Room 13-000, Toronto, Ontario, Canada M5G 2A2; Institute of Biomaterials and Biomedical Engineering, University of Toronto, 164 College Street– Room 407,Toronto, Ontario, Canada, M5S 3G9; School of Engineering, University of British Columbia, 1137 Alumni Ave, Kelowna, British Columbia, V1V 1V7, Canada; Faculty of Kinesiology and Physical Education, University of Toronto, 55 Harbord Street, Toronto, Ontario, Canada M5S 2W8; Department of Occupational Science and Occupational Therapy, University of Toronto, 500 University Avenue – Room 160, Toronto, Ontario, Canada M5G 1V7

**Keywords:** Balance recovery, handrail design, force generation, aging

## Abstract

Handrails have been shown to reduce the likelihood of falls. Despite common use, little is known about how handrail shape and size affect the forces that people can apply after balance loss, and how these forces and the corresponding ability to recover balance depend on age. Following rapid platform translations, 16 older adults and 16 sex-matched younger adults recovered their balance using seven handrail cross-sections varying in shape and size. Younger adults were able to withstand higher perturbations, but did not apply higher forces, than older adults. However, younger adults achieved their peak resultant force more quickly, which may reflect slower rates of force generation with older adults. Considering handrail design, the 38mm round handrails allowed participants to successfully recover from the largest perturbations and enabled the highest force generation. Conversely, tapered handrails had the poorest performance, resulting in the lowest force generation and withstood perturbation magnitudes. Our findings suggest that the handrail cross-sectional design affects the magnitude of force generation and may impact the success of recovery. Our findings can inform handrail design recommendations that support effective handrail use in demanding, balance recovery scenarios.

## Introduction

Falls on stairs cause over 1 million emergency room visits each year in the United States (Blazewick et al., 2018). Handrail use is a common strategy for recovering from balance loss (Gosine et al., 2021, 2019; King et al., 2011). Handrails can significantly reduce the risk of falls (Maki et al., 1998), provided that their design allows individuals to apply high forces to stabilize the body. Therefore, the ability to use the handrail to generate forces is an important aspect of handrail design.

An important but unanswered question is how the cross-sectional design of handrails affects the forces that young and older adults can apply when recovering balance. This information is important for informing standards such as Canadian building codes (Canada, 2015), which currently permit a wide range of handrail shapes and sizes. In the context of voluntary grasping, smaller handles (38mm to 51mm diameter) have been shown to permit higher grip forces than larger handles (76mm diameter) (Ayoub and Presti, 1971; Edgren et al., 2004; Pheasant and O’Neill, 1975). Similarly for handrails, 38mm to 44mm round handrails allowed the highest voluntary forces, while large decorative and rectangular handrails resulted in the lowest forces (Maki, 1985). By contrast, Dusenberry et al. reported that decorative handrails resulted in the highest forces, presumably because individuals could grip indentations on the side (Dusenberry et al., 2009). However, these results from voluntary grasping may not apply to reach-to-grasp reactions following balance disturbances, where the individual must first quickly and accurately contact the handrail, before applying adequate forces to stabilize the body (Komisar et al., 2019a).

In this study, we evaluate the effect of handrail cross-section on force generation during reach-to-grasp balance reactions. As strength decreases with age (Günther et al., 2008; Pasco et al., 2020), we were also interested in the effect of age on force generation and balance recovery. We considered forward and backward balance loss, which are common in stairway falls (Svanstrom, 1973). We hypothesized that 38mm round handrails would allow participants to withstand higher perturbation magnitudes and apply higher forces compared to larger handrails. We also hypothesized that younger adults would apply higher forces to the handrail, and would withstand higher perturbation magnitudes than older adults.

## Methods

### Participants

Healthy older (>65 years) and younger adults (18-35 years) were recruited from the community. Exclusion criteria included self-reporting of neurological, musculoskeletal or vestibular disorders that affect walking ability, heart disease or diagnosed osteoporosis. The study was approved by the institution’s research ethics board. All participants provided written informed consent.

### Experimental Environment

Data collection took place inside a 5mx5m laboratory mounted to a robotic platform (Figure 1). Participants stood next to a handrail, sloped at 38 degrees, at a height of 34” above the ground when in line with the participant’s toes. This corresponds to a staircase with a 7.75” rise and 10” run and the minimum handrail height allowable in the current National Building Code (Canada, 2015). The handrail was rigidly attached to a braced structure that allowed easy attachment of different handrail cross-sections. The handrail was instrumented with two tri-axial load cells (AMTI BP250500-2000, Advanced Medical Technology Inc., Watertown, MA) to measure applied forces.

**Figure 1:**
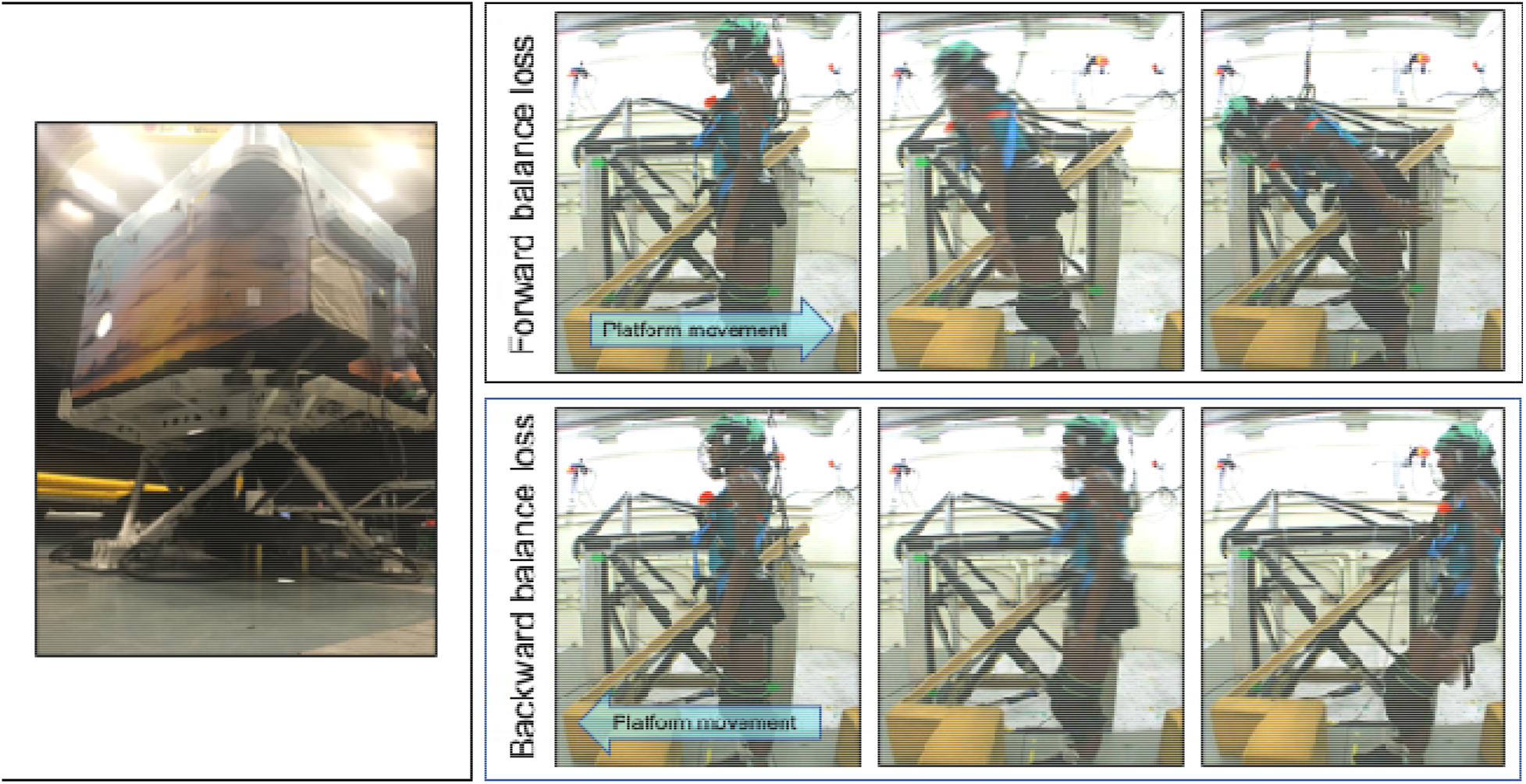
(Left) Laboratory mounted to robotic platform. (Right) Inside the lab. Snapshots of forward (top) and backward (bottom) balance loss, induced by backward and forward platform translation respectively.

We tested seven handrail cross-sections: 38mm (1.5”), 64mm (2.5”) and 76mm (3”) round, 64mm and 76mm “tapered”, and 64mm and 76mm “decorative” handrails (Figure 2). The decorative handrails were selected to represent common, commercially available handrails. To facilitate custom-sized fabrication and allow comparison between the shapes, the tapered handrails were simplified versions of commercially available “6010” milled handrails.

**Figure 2:**
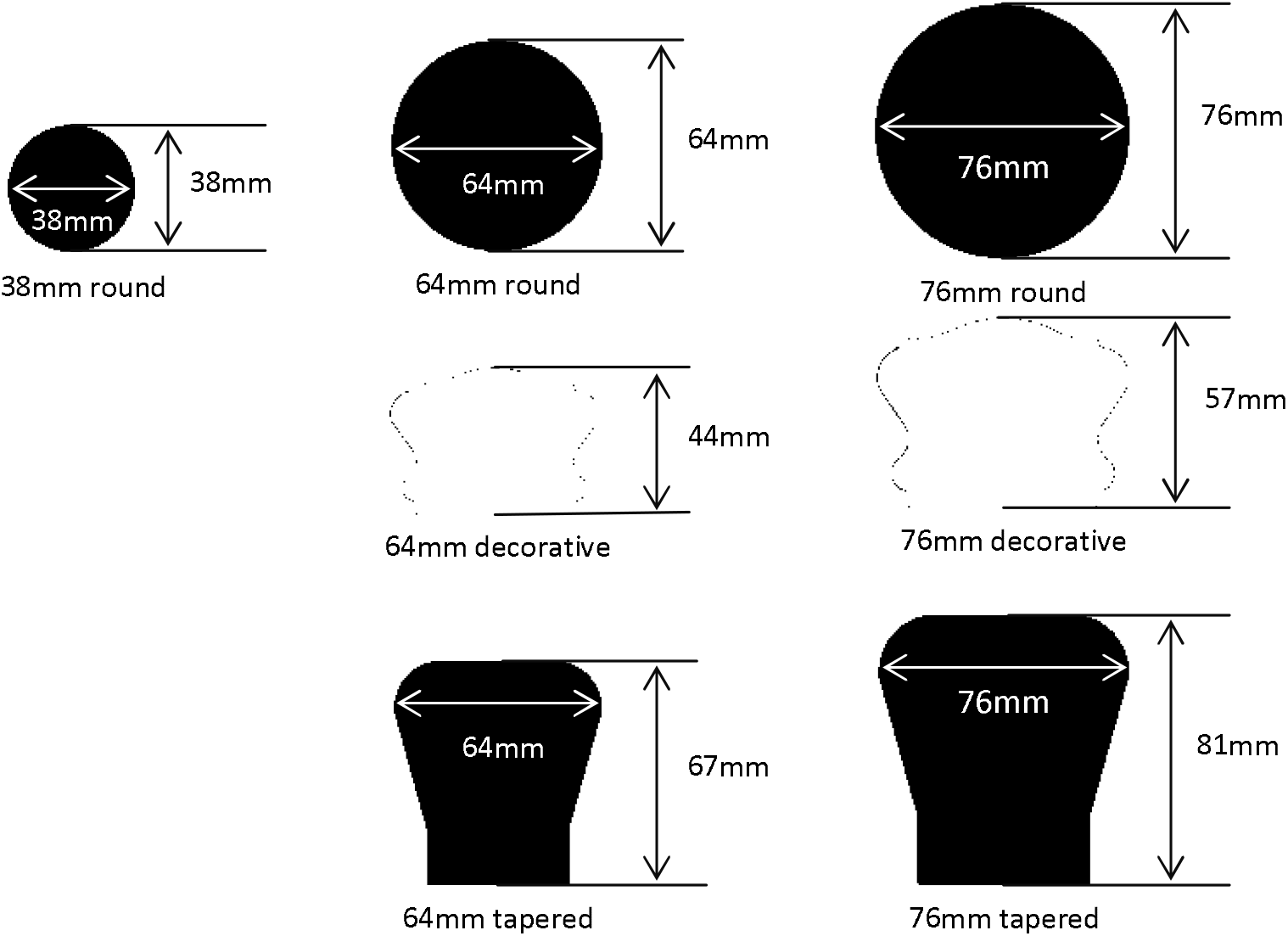
Depiction of the 38mm (1.5”), 64mm (2.5”) and 76mm (3”) round, 64mm and 76mm decorative, and 64mm and 76mm tapered handrails included in the study.

A fall arrest harness, knee pads, a hockey helmet, and an elbow pad on the right arm were used to reduce injury risk should balance recovery have failed. A load cell (AMTI MC3A-6-1000, Advanced Medical Technology Inc., Watertown, MA) in the harness line recorded the approximate support that the harness provided to the participant. Right-hand grip strength was tested using a Jamar handheld dynamometer (Jamar^®^ Hydraulic Hand Dynamometer; Patterson Medical, Mississauga, ON).

### Protocol

We focused on reach-to-grasp reactions because most younger adults and some older adults do not use the handrail proactively (Cohen and Cohen, 2001) and reach-to-grasp reactions tend to be more challenging for balance recovery. Participants stood in a comfortable position with their hands at their sides and their midline 0.6m away from a handrail (Gosine et al., 2019). They were instructed to recover their balance by grasping the handrail with their right hand, without stepping or relying on the harness. Foam blocks were placed around the participants’ feet to deter stepping. To reduce the effect of anticipation, participants counted backwards from a randomly selected number.

To disrupt balance, we used a “ramp-up” protocol of rapid platform translations with a square-wave acceleration profile (300ms acceleration phase followed by an equal and opposite 300ms deceleration phase). Participants experienced a small perturbation (1.0m/s^2^ for older adults and 1.5m/s^2^ for younger adults), which was increased in increments of 0.5 m/s^2^ to a maximum of 5m/s^2^ (Komisar et al., 2019b). The balance recovery was considered unsuccessful if the participant stepped; relied on the harness (defined by a harness load of >20% body weight (Carbonneau and Smeesters, 2014)); grasped the handrail with their opposing hand; or touched the floor, foam blocks or the supporting structure. Following an unsuccessful balance recovery, the perturbation magnitude was repeated until there were two unsuccessful trials (failure) or two successful trials (perturbation magnitude was increased). The timing and direction of the perturbations were randomized to reduce anticipatory movement.

To allow participants to become accustomed to the platform movement, participants completed a set of 12 familiarization trials using the ramp-up protocol (Cheng et al., 2012; Komisar et al., 2019b). Following the familiarization trials (not analyzed), the ramp-up perturbation protocol was completed for all handrail cross-sections, including again for the cross-section used for the familiarization trials. The order of testing handrail cross-sections was counter-balanced. Participants could rest for as long as needed between handrail cross-sections to reduce fatigue.

### Data analysis

We evaluated the effectiveness of the handrail by the ability to enable balance recovery, while supporting applied force in a variety of directions. For effectiveness, we considered the “Maximum Withstood Perturbation” (MWP). The MWP was defined as the trial completed at the highest perturbation magnitude that was successfully achieved prior to failure, just prior to the handrail no longer being effective (Komisar et al., 2019b). A higher MWP indicated that the handrail cross-section permitted recovery from larger destabilizations.

For each MWP trial, the peak resultant handrail force, and peak handrail forces in each Cartesian axis were determined in the first 1.5s after handrail contact. This time window captured the balance recovery response; inspection of the data confirmed peak applied forces occurred within this identified time frame. To determine the rate of force generation, time from handrail contact (resultant force > 10N) to peak resultant handrail force was measured. For all data analysis, applied handrail forces were filtered using a low-pass Butterworth filter of 10Hz.

### Statistical analysis

Age-related differences in participant demographics (height, weight, hand size, and grip strength) were determined using independent *t*-tests. Differences in self-reported handedness between age groups was determined using Fisher’s exact test.

Data were rank-transformed to correct for violations of normality and homogeneity of variance (Conover and Iman, 1981). General Linear Models were used to test the effect of handrail cross-section (within-subject factor) and age group (between-subject factor) on MWP. We also used General Linear Models to test the effect of handrail cross-section and age group peak forces and timing, with perturbation magnitude included as a co-variate. Participant ID was included as a random factor. Where significant main or interaction effects were found, post hoc pairwise comparisons with Tukey adjustments were performed to identify differences between handrail types. For force directions where perturbation magnitude was significant as a co-variate, we analyzed the correlation of peak force and perturbation magnitude. All statistical analysis was completed using SAS Enterprise Guide 7.1 (SAS, Cary, NC). Significance levels of *p*<0.05 were used for all outcome measures. Backward and forward balance loss were analyzed separately.

## Results

Sixteen older and 16 sex-matched younger adults were included in this analysis. We found no significant differences between groups for demographic data (Table 1).

**Table 1:** Demographic data *indicates a significant effect of age (p<0.05)

| Mean (SD) | Age (y) | Gender | Height (cm) | Weight (kg) | Hand length (cm) | Right hand grip strength (kg force) |
| --- | --- | --- | --- | --- | --- | --- |
| **Younger** | 23(3) | 10M, 6F | 171(8) | 73(15) | 18(1) | 39(9) |
| **Older** | 67(4) | 10M, 6F | 170(8) | 77(14) | 18(1) | 36(9) |

### Effect of handrail cross-section and age on ability to withstand perturbations

Younger adults withstood greater perturbation magnitudes than older adults, in forward and backward balance loss (*p*<0.003; Figure 3). Participants withstood higher perturbations with the 38mm round handrails than both tapered handrails (*p*≤0.003) in forward balance loss. In backward balance loss, the differences between handrails were not significant (*p*=0.249). There were no significant interactions between handrail design and age in either balance loss direction (p≥0.689).

**Figure 3:**
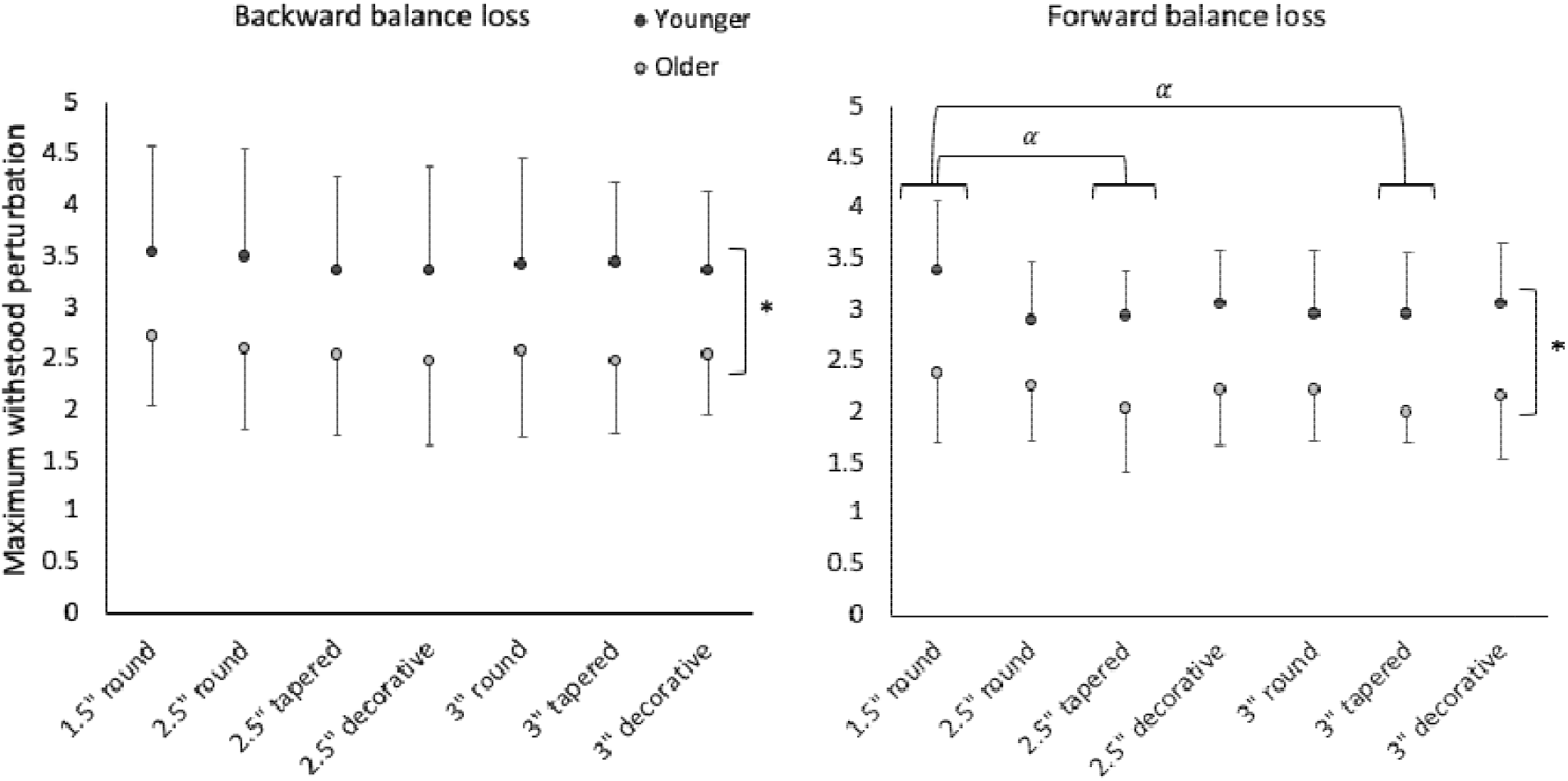
Maximum withstood perturbation for backward (left) and forward (right) balance loss *indicates a significant effect of age, α indicates a significant effect of handrail cross-section (p<0.05)

### Effect of handrail cross-section on applied handrail forces

#### Peak resultant handrail force

Participants applied mean peak forces to the handrail that ranged from 18.5% to 41.8% of bodyweight (Table 2, Figure 4A & 5A). As a covariate in analysis, perturbation magnitude significantly affected resultant handrail force for both forward and backward balance loss (*p*<0.001). For both balance loss directions, greater perturbation magnitudes were strongly correlated with greater peak resultant forces (|r|≥0.660, p<0.001).

**Table 2:**
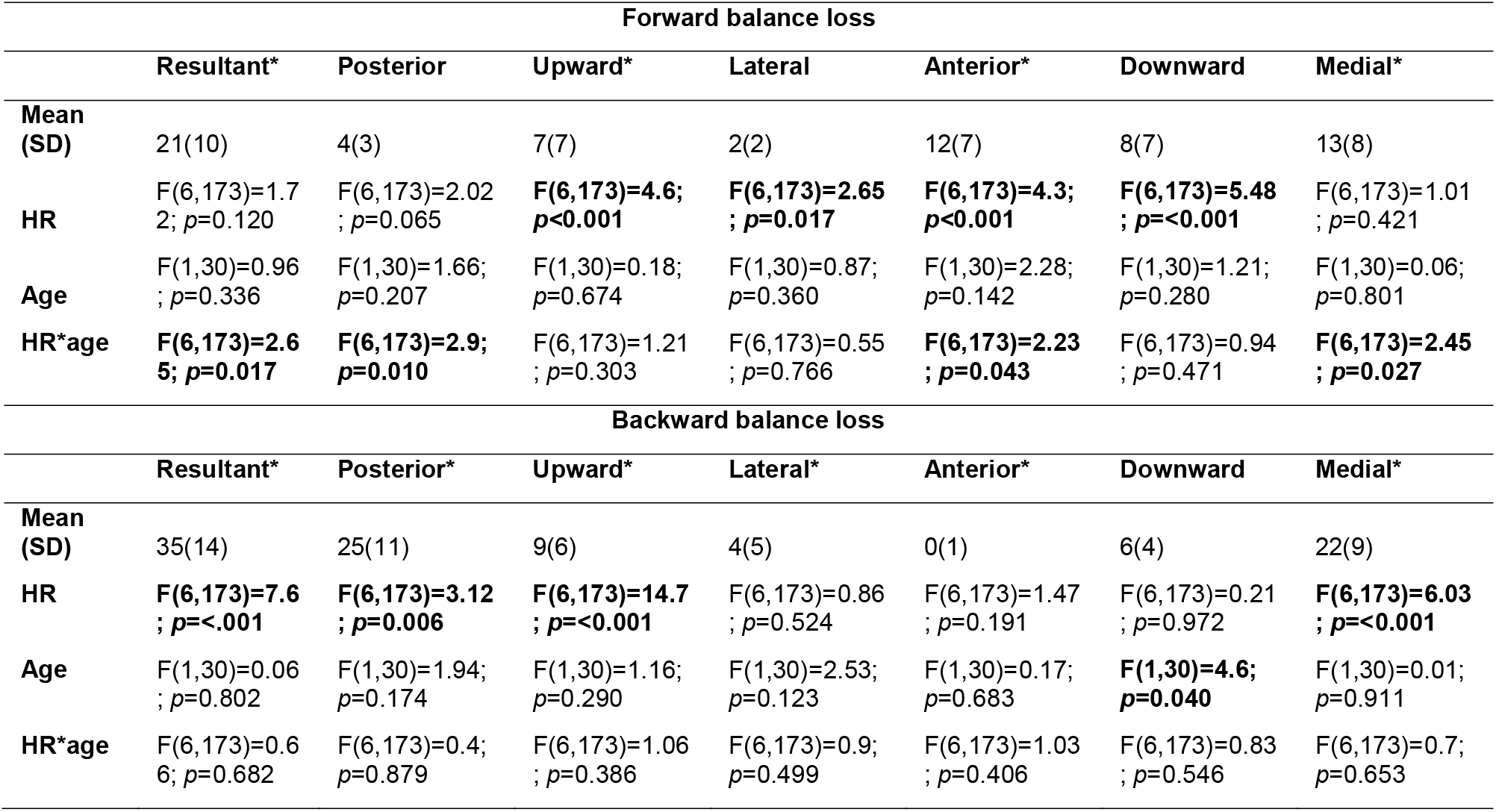
Overall mean and standard deviation of handrail forces (% bodyweight). Bolded text indicates a significant main or significant interaction effect. *indicates perturbation magnitude has a significant effect as co-variate. (p<0.05)

**Figure 4:**
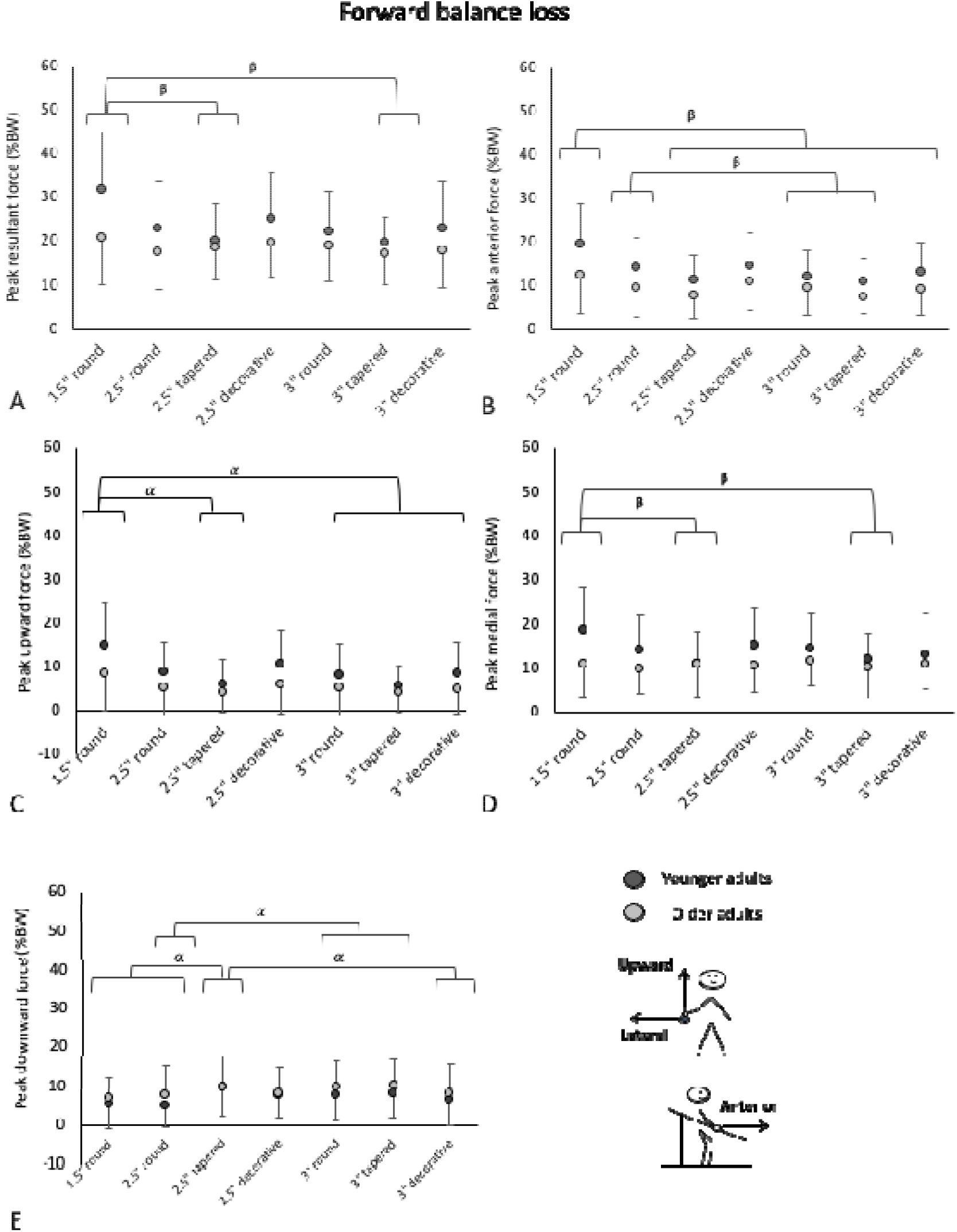
Mean handrail forces to recover from forward balance loss for younger (black circles) and older adults (grey circles), across the various handrail cross-sections. The applied force directions (% body weight, %BW) where a significant effect of handrail cross-section was found are presented in the figure. Peak resultant force [A]; peak anterior force [B]; peak upward force [C]; peak medial force [D]; peak downward force [E]. α indicates a significant main effect of handrail cross-section (p<0.05), β indicates a significant effect of handrail cross-section for young adults only (p<0.05)

In forward balance loss, a significant interaction of age and handrail cross-section on peak resultant force was identified (*p*=0.018). For older adults, handrail cross-section did not affect resultant force; for younger adults, the 38mm round handrail resulted in higher resultant forces than both tapered handrails (*p*≤0.026).

In backward balance loss, handrail cross-section affected peak resultant force (p<0.001). The 38mm round handrail resulted in higher peak resultant forces than all handrails except the 64mm round rail (*p*≤0.001). The 64mm round handrail also resulted in higher forces than the 76mm tapered handrail (*p*=0.043).

#### Peak handrail forces in cartesian coordinates

Participants applied forces between 0% to 30% of bodyweight across the different axes (Table 2). Forces were primarily applied in the medial direction and the direction of balance loss (i.e. posterior force in backward balance loss and anterior force in forward balance loss). Differences between handrail cross-sections and age are also presented in Figures 4 & 5, for directions of force considered to have functionally meaningful differences (defined as difference in mean peak forces greater than 1.25% BW (Maki et al., 1998)).

##### Forward balance loss

Participants applied the highest overall mean forces in the anterior and medial directions, with large contributions in the upward and downward directions (Figure 4). The smallest forces were in the lateral direction. Perturbation magnitude was a significant covariate for upward, anterior and medial directions of applied force (*p*≤0.001), reflecting greater forces with larger perturbations (|r|> 0.498, p<0.001).

Handrail cross-section affected the peak upward and downward forces (*p*≤0.017) across both groups. 38mm handrail resulted in greater upward force than the 64mm tapered handrails and all 76mm handrails (*p*≤0.047). The 64mm tapered handrails resulted in higher downward forces than the 38mm round, 64mm round and 76mm decorative handrails (*p*≤0.050). The 76mm round and tapered handrails resulted in larger downward forces than the 64mm round handrail (*p*≤0.023).

Age and cross-section interacted significantly for the peak posterior, anterior and medial forces (*p*≤0.043). When separated by age group, handrail cross-section affected peak posterior force for older adults (older: p=0.028, younger: *p*=0.077); however after Tukey corrections, pairwise comparisons of handrail cross-section were not significant (*p*≥0.057). For anterior and medial forces, while there was an effect of handrail cross-section for younger adults (*p*≤0.009), but not for older adults (*p*≥0.096). In the anterior direction, 38mm and 64mm round handrails had higher forces than the 76mm round and 76mm tapered handrails and the 38mm round handrails had higher forces than the 64mm tapered, 64mm decorative and 76mm decorative handrails (*p*≤0.037). In the medial direction, the 38mm round handrail had higher forces than the 64mm tapered and the 76mm decorative handrails (*p*≤0.030).

Age did not affect applied handrail forces in any direction (*p*≥0.142).

##### Backward balance loss

Participants mostly applied forces in the posterior and medial directions, with smaller contributions in the upward, lateral and downward directions (Figure 5). Forces in the anterior direction were negligible (mean <1% BW). As a covariate, perturbation magnitude was significant for posterior, upward, lateral, anterior and medial directions of applied force (*p*≤0.001), reflecting greater forces with larger perturbations (|r|>0.324, p<0.001).

**Figure 5:**
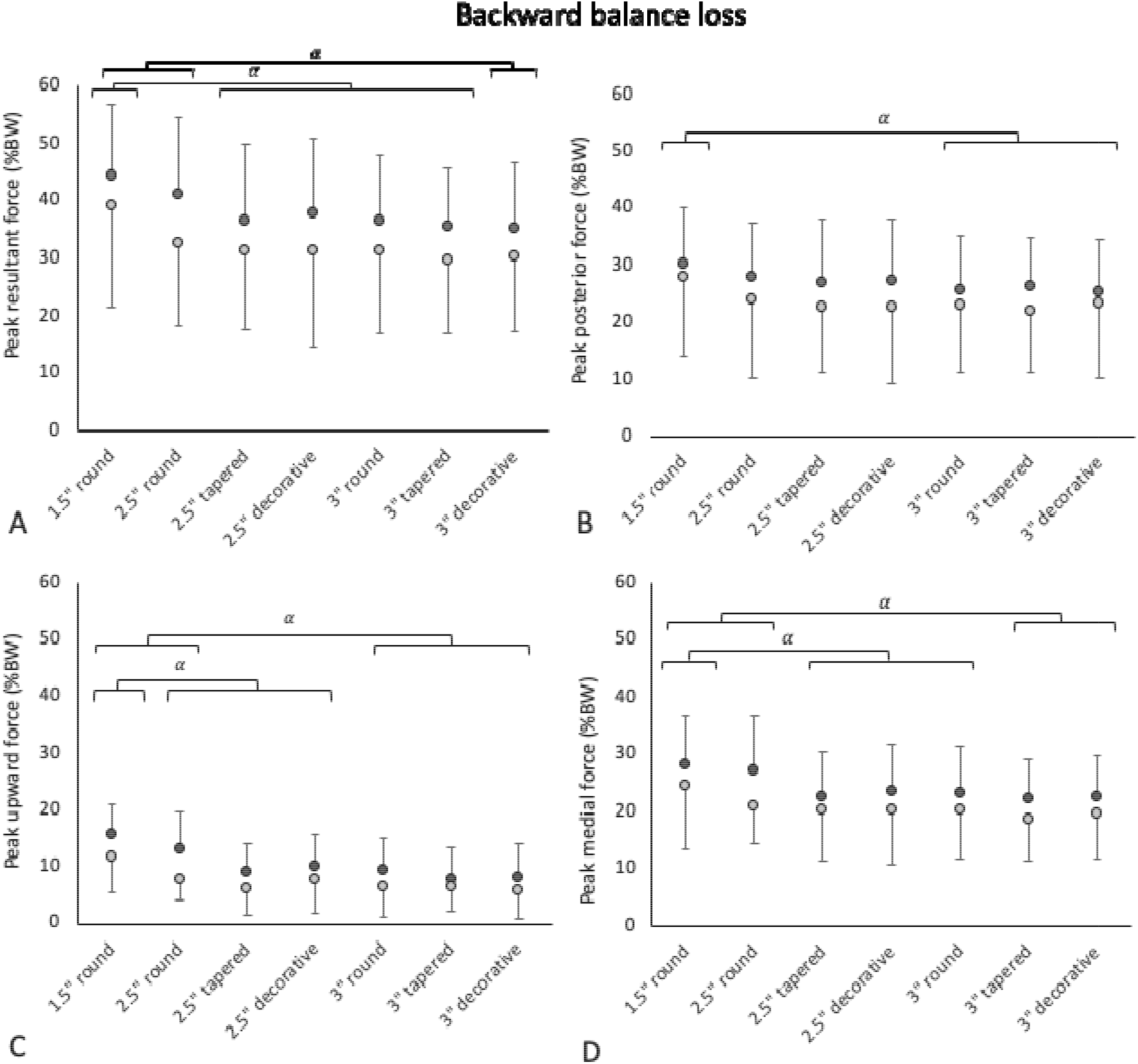
Mean handrail forces to recover from backward balance loss for younger (black circles) and older adults (grey circles), across the various handrail cross-sections. The applied force directions (% body weight, %BW) where a significant effect of handrail cross-section was found are presented in the figure. Peak resultant force [A]; peak anterior force [B]; peak upward force [C]; peak medial force [D]. α indicates a significant main effect of handrail cross-section (p<0.05), β indicates a significant effect of handrail cross-section for young adults only (p<0.05)

Higher upward forces were seen with the 38mm round handrail than all handrails (*p*≤0.019). Furthermore, the 64mm round handrail resulted in higher upward forces than all the 76mm handrails (*p*≤0.07). In the medial direction, the 38mm round handrail resulted in higher forces than all handrails except the 64mm round handrail (*p*≤0.006). The 64mm round handrail resulted in higher forces than the 76mm decorative and both sizes of tapered handrails (*p*≤0.046).

Older adults applied greater downward forces than younger adults (*p*=0.016). No other significant effects of age were detected for forces in other directions (*p*≥0.057).

#### Timing of peak applied resultant force

Perturbation magnitude was not a significant covariate for timing variables for either balance loss direction (*p*≥0.238). Older adults had slower time from handrail contact to peak resultant handrail force than younger adults in backward balance loss (271ms vs 205ms; *p*=0.001) but not forward balance loss (380ms vs 262ms; *p*=0.130). There was also an effect of handrail cross-section on timing of applied resultant force following backward balance loss (*p*=0.003), but not forward balance loss (*p*=0.545). In backward balance loss, the 64mm tapered handrails had slower time to peak force than the 64mm decorative and 76mm round handrails (*p*<0.020).

## Discussion

Handrails should be designed such that individuals can apply high forces to recover from balance loss and reduce the risk of injury. Many different handrail designs exist in homes, public buildings and outdoor spaces (Canada, 2015), and the shape and size of handrails may affect one’s ability to apply high forces for balance recovery (Maki, 1985). This study was the first to evaluate the effect of handrail cross-section design on the ability of young and older adults to withstand balance loss, and on the concomitant applied handrail forces.

### Effect of handrail cross-section on balance control and force generation

As hypothesized, we found that use of the 38mm handrails involved the highest applied handrail forces in both balance loss directions. The 38mm handrails were also associated with the highest perturbation magnitudes during forward balance loss, particularly compared to the tapered rails. These findings align with literature that reported that 38mm round handles resulted in the highest applied forces (Edgren et al., 2004; Grant et al., 1992), the longest duration of force generation before onset of fatigue (Ayoub and Presti, 1971), and the highest force-to-effort ratio (Ayoub and Presti, 1971). In the context of handrails, Maki (1985) demonstrated that 38mm handrails also allowed generation of the highest voluntary forces compared to larger cross-sections and different shapes. Our results extend this work to show that 38mm round handrails enabled higher handrail forces while recovering from larger perturbations. Although our task differs from stair gait, it would follow that in the real world, handrail designs that enable recovery from larger destabilizations would in turn support recovery from a greater range of destabilizations and result in fewer falls.

In contrast to the 38mm round handrails, the tapered handrails resulted in the smallest withstood perturbations in forward balance loss. In general, the tapered handrails also resulted in the lowest resultant, medial, upward and anterior forces but the highest downward forces. The findings are not surprising since the tapered handrails had the largest cross-sectional area, which impedes the ability to use a power grip as the fingers cannot wrap around the underside of the handrail. In line with our findings, literature has shown that handhold shapes and sizes that facilitate a power grip enabled much higher forces to be generated than designs that require a pinch grip (such as a rectangular handrail) (Maki, 1985; Swanson, 1970). With the tapered handrails, participants may have also compensated for low anterior and upward forces by pushing harder in the downward direction. The reliance on downward forces to assist with balance recovery may be less effective during ongoing stair descent, where participants have been seen to pull along the handrail to recover their balance (Gosine et al., 2021, 2019; Maki et al., 1998). These findings also highlight the importance of evaluating handrails during balance recovery, where forces are applied in multiple directions.

Our results shed light on balance recovery using handrails with complex geometry. For example, Dusenberry et al. (2009) has suggested that the indentation of tapered handrails enhanced the ability to grip the handrail, permitting comparable force generation in the medial and anterior-posterior directions to that of a 2” round handrails. However, Dusenberry et al. (2009) restricted the use of the underside of the handrail. While the extent of tapering in our current study may have differed from that of Dusenberry et al. (2009), our findings suggest that a handrail which permits at least some grasping of the underside of the handrail during reactive grasping is important to facilitate force generation. While the 64mm decorative handrails did not perform as well as the 38mm round handrails, participants were able to generate similar resultant, upward and medial forces in forward balance loss, perhaps because of the size of the 64mm decorative handrail rather than the shape itself. For the 64mm decorative handrails, participants may have used the underside of the handrail allowing for larger force application. Further analysis of applied forces associated with varying grip types is needed to generalize these results to handrails beyond the seven tested in this study, and to understand key design features that enable high force generation in multiple directions.

Unlike forward balance loss, we found no significant effect of handrail cross-section on perturbation magnitude during backward balance loss. This may be because participants could apply higher forces and withstand higher perturbations across all handrails in backward balance loss. The angle of the handrail allowed participants to lean against the handrail when recovering from backward balance loss. Hence, the shape and size of the handrail may have been less important for balance recovery. Alternatively, participants may have gripped the rail at a higher position when falling backward. We previously reported that young adults could withstand higher perturbations with higher handrails (Komisar et al., 2019a). This highlights the importance of other handrail design characteristics in addition to cross-section, such as height, to support safe balance recovery.

### Effect of age on force generation and balance control

Despite experiencing significantly lower perturbation magnitudes across all handrail cross-sections, older adults applied similar forces to younger adults. One possible explanation is that older adults relied more heavily on external stabilizing moments instead of internal stabilizing strategies (e.g., “hip” or “ankle” strategies), potentially due to reduced muscle strength (Pasco, et al., 2020). Our findings align with other evaluations of balance recovery (where stepping was permitted), where older adults relied more on handrail grasping rather than using internal stabilizing strategies, and took more compensatory steps (Maki et al., 2000; McIlroy and Maki, 1996). Further research with surface electromyography is needed to identify the role of internal stabilizing strategies in balance recovery when handrails are available, and how these strategies vary with age.

We also found that older adults took longer to reach their peak force after handrail contact when compared to younger adults. This may reflect slower rates of force generation, which has been consistently demonstrated for other muscle groups (Mackey and Robinovitch, 2006; Watanabe et al., 2011). While both young and older adults applied similar peak handrail forces, the failure of older adults to quickly apply their peak forces after contact may have contributed to difficulties in controlling their COM momentum – resulting in lower MWPs the younger adults. The slower rate of handrail force generation in older adults highlights the importance of proactive rail contact, to reduce the overall time to achieve peak handrail force relative to balance loss (McKay et al., 2013).

We acknowledge the study limitations. First, participants stood a fixed distance away from the handrail (60cm). While the selected distance allows most of the population to reach the handrail (Gordon et al., 1989), this may be further away than some people would typically walk (Cohen, 2000; Cohen and Cohen, 2001). Standing closer to the handrail has been associated with smaller medial forces during balance recovery (Maki et al., 1998), and further work should explore different initial stance conditions when evaluating handrail cross-sectional design. Second, our protocol involved perturbations of upright stance, where participants were instructed not to step. Handrail loading characteristics may differ in contexts where participants can also step following balance loss, such as during stair descent (Gosine et al., 2021, 2019). Future studies should consider forces applied to different handrail designs when recovering from unexpected balance loss during ongoing stair descent. Finally, our study focused on healthy adults. Hand impairments in older adults are common and affect grip strength (e.g., arthritis (Zhang et al., 2002)); however, these individuals were not included in this work. Further research is needed to understand the effect of handrail shape and size on balance recovery in these populations.

We analyzed the effect of handrail cross-section on the ability to recover balance and generate forces throughout recovery. Despite younger adults withstanding higher perturbations, both groups generated similar peak forces. Small, round handrails were the most effective handrails tested, as they allowed participants to successfully recover from the largest perturbations while applying the highest forces in most cases, while tapered handrails were the least effective. Designing handrails that enable high force generation to facilitate successful balance recovery for both younger and older adults is an important step towards safer stairway design.

## Conflict of interest

None.

## Acknowledgements

The authors would like to acknowledge financial support from the Canadian Institutes of Health Research Operating Grant (CIHR MOP 142178), NSERC Canadian Graduate Scholarships-Doctoral Award (P Gosine), AGE-WELL Network of Centres of Excellence Postdoctoral Awards in Technology and Aging (V Komisar), and a MITACS Accelerate Award (V Komisar). The authors thank Dr. Bruce Haycock, Ms Susan Gorski, Mr Roger Montgomery and Mr Dan Smyth for their technical assistance and support.

## Notes

### Competing Interest Statement

The authors have declared no competing interest.

